# Reduced blood to brain glucose transport as the cause for hyperglycemia in a model resolves multiple anomalies in type 2 diabetes

**DOI:** 10.1101/2022.01.19.477014

**Authors:** Akanksha Ojha, Milind Watve

**Affiliations:** Indian Institute of Science Education and Research, Pune, Pune, 411008, India; Independent Researcher, E-1-8, Girija Shankar Vihar, Karve Nagar, Pune 411052, India

**Keywords:** Glucose homeostasis, hyperglycemia, type 2 diabetes, steady state

## Abstract

Classically type 2 diabetes is believed to be a result of insulin resistance and relative insulin deficiency. However, evidences have been accumulating against the insulin resistance centered models. Absence of fasting hyperglycemia by insulin receptor knockouts or insulin suppression, evidence for hyperinsulinemia preceding insulin resistance, the perplexing hyperinsulinemic normoglycemic state, reduced glucose transport to the brain preceding hyperglycemia, signs of vasculopathy preceding hyperglycemia, absent or poor correlation between fasting glucose and insulin, very strong positive correlation between indices of insulin resistance and β cell function in population data are some of the anomalous findings which glucose homeostasis models have not addressed so far. With increasing evidence for neuronal involvement in glucose regulation, we propose a refined model of glucose regulation that considers brain glucose and insulin levels as the ultimate target of homeostasis and combines central and peripheral mechanisms of regulation. A model considering reduced rate of blood to brain transportation of glucose and insulin as primary pathology explains most of the patterns, with or without insulin resistance. Apart from resolving multiple anomalies the model also accounts for the limited and inconsistent success of glucose normalization in effectively reducing diabetic complications and mortality.

## 1. Introduction

Presently, any metabolic disorder with consistent raised plasma glucose and insulin resistance, which cannot be categorized into either type 1 diabetes or gestational diabetes is defined as type 2 diabetes mellitus (T2DM) (DeFronzo et al., 2015). It is the most common form of diabetes, accounting for over 90% of all cases. The currently perceived cause of hyperglycemia in T2DM is insulin resistance followed by failure of compensatory insulin response. Historically the role of brain in glucose homeostasis was revealed by experiments in which damage to the fourth cerebral ventricle was shown to impair homeostatic control (Bernard, 1859). However, at a later stage, pancreatic extracts were shown to lower glucose and eventually insulin was identified as the active principle (Banting et al., 1922). It is necessary to note that the differentiation of type 1 and type 2 diabetes was not made at that time. The success of insulin was so spectacular that all other mechanisms of glucose regulation were almost forgotten for about a century(Lundqvist et al., 2019). The distinction between type 1 and type 2 diabetes was to become clear half a century later and it was realized that unlike in type 1, insulin deficiency was not the primary cause of hyperglycemia in type 2 diabetes. However, by this time the thinking in the field was so much insulin centered that the inability of normal or increased levels of insulin to regulate plasma glucose was termed insulin resistance, without carefully evaluating alternative hypotheses.

Further, the insulin resistance hypothesis is criticized for having a circular logic which makes it un-falsifiable(Diwekar-Joshi&Watve, 2020). Insulin resistance is said to be responsible for the inability of normal or raised levels of insulin to regulate glucose. However, insulin resistance itself is measured by the inability of insulin to regulate glucose. There is no independent measure of insulin resistance that can be used to test the causal role of insulin resistance. Impairment of insulin signaling in experiments using (i) tissue specific insulin receptor knockouts(Blüher et al., 2002; Kadowaki, 2000) (ii) insulin degrading enzyme knockouts(Costes & Butler, 2014; Maianti et al., 2014)(iii) pharmaceutical insulin suppression by diazoxide and octreotide(Giustina et al., 1991; Lamberts & Hofland, 2019; Leahy et al., 1994; Matsuda et al., 2002)(iv) insulin gene suppression by RNAi (Mehran et al., 2012b)(v) alteration in insulin gene dosage(Templeman et al., 2015)does not consistently alter fasting glucose although it alters the post meal glucose curve. In epidemiological data, fasting insulin and fasting glucose are poorly correlated while post meal insulin and glucose are strongly correlated, unlike what would be expected by any of the classical models. With converging multiple lines of evidence Diwekar-Joshi and Watve (2020) raised doubts on whether insulin resistance and failure of compensatory hyperinsulinemia is a necessary and sufficient explanation of fasting hyperglycemia.

Many mathematical models have been constructed based on the assumption of insulin resistance. A common set of assumptions is shared by most of the models that fasting glucose is the steady state achieved by a balance between glucose uptake by tissues and glucose production by liver. One of the foundational assumptions is that both the above processes are regulated by insulin signaling in the steady state(Matthews et al., 1985). Therefore, insulin signaling has been central to these models. Since recent research has exposed the inadequacies of this model (Diwekar-Joshi & Watve, 2020)there is a need to hunt for better alternative models.

An alternative to the classical model which assumes brain glucose level as the target of homeostasis has been suggested and mathematical models incorporating this concept have been constructed(Gaohua & Kimura, 2009; Watve, 2013). These models have not been explored sufficiently towards a comparative evaluation with peripheral models with respect to their predictions and their physiological as well as clinical implications. Here we refine the brain centered model further incorporating recent evidence, evaluate its performance and explore the physiological and clinical implications.

### 1.1 What a glucose homeostasis model needs to explain

Most models of glucose homeostasis have a limited goal of explaining altered fasting steady state glucose level and the alterations in the glucose curve in the pre-diabetic and diabetic state. However, now we have a large number of experimental results that are potentially anomalous. A model prediction matrix, in which each model is tested for matching its prediction with multiple known empirical patterns, is an appropriate and robust approach to compare alternative models. Such an approach has been used to test alternative hypothesis in a number of other contexts(Shinde et al., 2021; Thakar et al., 2003; Vibishan & Watve, 2020; Watve & Diwekar-Joshi, 2016) and it can be the right approach to test alternative models of glucose homeostasis in T2DM. A model of choice would be one which can predict all or most of the known physiological patterns in type 2 diabetes so that no or minimal anomalies are left. We take a few steps towards a model prediction matrix, although a lot more exploration of many other models is required before constructing such a matrix.

Following is the list of known empirical patterns that a model needs to be compatible with.

1. The model should allow stable steady state glucose on fasting: Fasting glucose is assumed to be a steady state with adequate evidence(Lerner & Porte, 1972; Matthews et al., 1985; R. C. Turner et al., 1979). Achieving a steady state is one of the primary objectives of every homeostasis model. Many of the key concepts of classical models are also based on achieving a steady state in the fasting condition(Matthews et al., 1985; R. C. Turner et al., 1979).
2. The model should also allow stable steady state insulin on fasting. Normally this is not a problem for any model, however many models have a problem in explaining the altered steady state insulin level in a prediabetic state as explained below.
3. Prediabetic state of normoglycemia with hyperinsulinemia: A prediabetic state is often accompanied by high level of fasting insulin (*FI*) but normal fasting glucose (*FG*) in the plasma. This is not easy to achieve in a model. The classical qualitative explanation of this state is that it is a state of insulin resistance with compensatory rise in *FI* such that *FG* remains normal. For this to happen, there is a need to estimate the level of insulin resistance and regulate insulin secretion accordingly. It is possible to hypothesize a mechanism of sensing insulin resistance and conveying this information to the pancreas. Such a mechanism can be glucose dependent or glucose independent. Currently no glucose independent mechanism of accurately assessing the insulin resistance and conveying it to β cells is known. A common assumption is that insulin resistance reduces glucose uptake thereby increasing *FG*. The rise in *FG* stimulates insulin production and the rise in insulin levels normalizes glucose again. However, this mechanism fails to achieve a steady state hyperinsulinemic normoglycemic condition(Diwekar-Joshi & Watve, 2020). Plasma insulin has a short half-life and after glucose levels are normalized, it is unlikely to stay at a higher level unless there is a glucose independent trigger for insulin secretion. Explaining a hyperinsulinemic normoglycemic steady state is a tricky challenge that any model needs to meet.
4. Hyperinsulinemia precedes insulin resistance and T2DM: Although hyperinsulinemia is believed to be a compensatory response to insulin resistance, a number of studies show that hyperinsulinemia precedes obesity and insulin resistance (Corkey &Shirihai, 2012; Mehran et al., 2012a; Pories & Dohm, 2020; Shanik et al., 2008; Weyer et al., 2000; Wiebe et al., 2021). This raises two independent questions. One is how and why hyperinsulinemia precedes insulin resistance. The second, more perplexing question is that if hyperinsulinemia is not a compensatory response to insulin resistance, the failure of compensation explanation becomes redundant. This needs to be replaced by a more coherent explanation of hyperglycemia.
5. The model should explain the following features of post meal glucose curve and its alterations in prediabetic and diabetic states, which cumulatively are called impaired glucose tolerance curve.

a. Increased height of peaks
b. Delayed return to normal
c. Increase in the time difference in glucose and insulin peaks in the diabetic state.
6. The normal fasting glucose with impaired glucose tolerance (NFG-IGT) state: It is common to find that in a prediabetic state the glucose tolerance curve is altered with normal level of fasting glucose.
7. Stress hyperglycemia: Why and how stress causes hyperglycemia only in some individuals (Dungan et al., 2009) needs to be explained by a model with realistic and testable assumption.
8. Hyperglycemia after intensive exercise: If in a fasting state *FG* is a resultant of the rate of liver glucose production and glucose disposal, exercise that increases glucose disposal should decrease *FG*. However, in some studies plasma glucose remains unchanged or is increased after exercise (Coggan, 1991; Marliss & Vranic, 2002). A model needs to incorporate this phenomenon.
9. An important challenge is to explain why insulin receptor knockouts specific to muscle, fat cell or liver results into normal fasting glucose but altered post glucose load curves. Similarly, insulin degrading enzyme knockouts do not alter *FG*. Also insulin suppression by agents such as octreotide or diazoxide fail to alter *FG*(Diwekar-Joshi & Watve, 2020). A model should explain this observation. Diwekar-Joshi and Watve (2020) further differentiated between the consequential steady state (CSS) and targeted steady state (TSS) models and demonstrated that the results of the insulin receptor knock out and insulin suppression experiments can be explained by a TSS but not by a CSS model. In effect, to be able to account for these results a model needs to be a TSS model.
10. Reduced glucose transport to brain, particularly in obese or prediabetic individuals: It is known over a long time that in type 2 diabetes the rate of glucose transport from blood to brain is slowed down substantially(Gjedde & Crone, 1981). In rodent models, the glut-1 expression in brain capillaries is shown to be reduced(Cornford et al., 1995; Matthews et al., 1985; Pardridge et al., 1990). This has been viewed as a response to hyperglycemia. However, Hwang et al (2017) show that subnormal transport is evident in obese and presumably prediabetic state, even before hyperglycemia appears. A model needs to account for this pattern as a causal or consequential phenomenon.
11. The T2DM specific islet amyloid deposition which often accompanies islet deterioration: Islet amyloid deposition is frequently associated with type 2 diabetes but not observed in type 1. The causes of islet amyloid deposition should be compatible with the glucose homeostasis model (Höppener et al., 1999; Porte & Kahn, 1989; Watve, 2014).
12. B cell deterioration pattern: The deterioration of β cells appears to follow a peculiar pattern in T2DM, in which a substantial proportion of β cells survive lifelong (Bacha et al., 2013; Butler et al., 2003; Clark et al., 2001; Porte & Kahn, 2001; Watve, 2014). If dysfunction and damage to the β cell population is assumed to be an essential prerequisite of hyperglycemia in a model, the models need to account for the peculiar population dynamics of β cells.
13. Increased liver glucose production as well as ketogenesis with SGLT2 inhibitors: Over the last decade SGLT2 inhibitors, which allow greater clearance of glucose through urine, have offered a novel means of combating hyperglycemia. Lowering of plasma glucose by SGLT2 inhibitors is accompanied by increased liver glucose production as well as increased ketogenesis(Limenta et al., 2019; Mistry & Eschler, 2021; Op den Kamp et al., 2021; Pfützner et al., 2017).
14. The Somogyi phenomenon i.e., hyperglycemia following infusion of insulin is seen in at least some cases of T2DM but not in healthy individuals. This can happen without being overtly hypoglycemic (Campbell, 1976; Reyhanoglu & Rehman, 2021).
15. In cases of brain injury hyperglycemia is common (Shi et al., 2016). In bacterial meningitis hyperglycemia is often observed accompanied by low CSF glucose level(Krishnan et al., 2016; Schut et al., 2009). A model should be able to account for the apparent contradiction.
16. In population data one sees good correlation between post meal glucose and insulin but poor correlation between *FG* and *FI* (Diwekar-Joshi & Watve, 2020).
17. Although FG and FI are poorly correlated in prediabetic state, HOMA-β and HOMA-IR have strong positive correlation in population data (Diwekar-Joshi & Watve, 2020).
18. Failure of glucose normalization to reduce the frequency of complications and mortality: Unlike T1DM, in T2DM tight regulation of plasma glucose has failed to show reduction in mortality consistently across multiple large scale clinical trials (Diabetes Prevention Program Research Group, 2015; Ferrannini & DeFronzo, 2015; King et al., 1999; Klein, 2010; Lee et al., 2021; Schwartz & Meinert, 2004; R. Turner et al., 1998; The NICE-SUGAR Study Investigators, 2009, UK Prospective Diabetes Study (UKPDS) Group, 1998). If hyperglycemia is the primary pathology of type 2 diabetes, preventing or correcting it should have reduced the frequency of complications and mortality considerably.
19. Reversal of hyperglycemia by FGF 21 in all models of rodent diabetes: Independent of the cause of hyperglycemia, a single injection of FGF 21 was able to achieve long term normalization of plasma glucose (Laeger et al., 2017). This is a potential anomaly that a model needs to resolve.

While most models intend to explain 1, 2 and 5 of the above, they either fail to explain or do not address the others. Our expectation from a satisfactory model is that it should inherently predict these outcomes or should be compatible with them. If an additional explanation that does not contradict the model, accounts for the phenomenon, the model can be said to be compatible. Compatibility with empirical findings cannot be taken as a proof or validation of the model, but if a model directly contradicts one or more of the empirical findings, it becomes serious grounds for rejecting the model. Later when we describe model results, we will use the numbers in the above list in square brackets to indicate that the model accounted for this pattern.

### 1.2 Peripheral models of glucose homoeostasis

There is a rich history of development of mathematical models of glucose homeostasis and the origin and progression of type 1 and type 2 diabetes (reviewed by Ajmera et al., 2013; Mari et al., 2020). The focus of the field is so much on peripheral mechanism that the review by Mari et al (2020) does not even cite the models incorporating the role of brain. Ajmera et al (2013) briefly mention the Gouhua et al (2009) model but do not elaborate on its potential implications. A class of models attempts to capture glucose homeostasis at the systems levels whereas other models look at individual components such as insulin dependent glucose uptake, liver glucose production, β cell dysfunction, glucose stimulated insulin secretion and incretin effects at greater details. Nevertheless, these models together have not accounted for many of the patterns listed above. The central assumption of these models is more or less invariant and revolves around insulin resistance and the failure of insulin response to adequately compensate for it (Mari et al., 2020). Diwekar-Joshi and Watve (2020) claimed that any model with this set of assumptions is not compatible with empirical patterns 3, 4, 6, 9, 16 and 17 of the above. Whether any variation of these models can do so has not been adequately explored. Furthermore, all the peripheral regulation models are CSS models in that the fasting steady state is a consequence of the rate of liver glucose production and a concentration dependent rate of glucose uptake. In these models a change in either or both the rates inevitably alters the steady state. This contrasts the TSS models in which alteration in these rates alters the time required to reach a steady state but does not alter the steady state glucose level(Diwekar-Joshi & Watve, 2020). An attempt to develop a TSS model with only peripheral mechanisms has not been made to the best of our knowledge.

We do not intend to review glucose homeostasis models here. However, we observe from published reviews of glucose homeostasis models (Ajmera et al., 2013; Mari et al., 2020)that most models do not address the apparently anomalous empirical findings particularly 3, 4, 6, 9, 10, 11, 12, 13 and 15-18 among the ones listed above. A detailed exploration in to whether some variations of these models can address the apparently anomalous patterns is beyond the scope of this paper but we remain open to this possibility.

### 1.3 Brain centered models of glucose homeostasis

It is quite well known that a number of neuronal mechanisms in the brain are involved in energy homeostasis. Nevertheless, for some reason, they do not occupy a central stage in the mainstream thinking in T2DM and glucose regulation models. A number of recent publications highlight the role of brain in different ways(Lam, 2005; Osundiji et al., 2012; Perrin et al., 2004; Watve, 2013, Lundqvist et al 2019). Peters (2004), Gaohua and Kimura (2009) and Watve (2013) proposed that the primary target of glucose homeostasis is to regulate glucose levels in the brain. Plasma glucose levels are only a means to achieve required supply of glucose mainly to the brain. Since transport of glucose to the brain is more restricted, when adequate supply of glucose to the brain in ensured, it is likely that the supply to other organs already gets ensured.

In the Gaohua and Kimura (2009) as well as in the Watve (2013) model the rate of glucose transport across the BBB is assumed to be an adaptation to hyperglycemia. The finding that glucose transport is reduced to an intermediate level in obese and prediabetic individuals (Büsing et al., 2013; Hwang et al., 2017)suggests that the deficiency in transport precedes hyperglycemia rather than following it. Therefore, it is more likely to be causal than consequential. Glucose deficiency in the brain is known to induce liver glucose production and suppresses insulin release through autonomic control(Boland et al., 2017). Sympathetic tone has also been shown to be higher in T2DM (Thackeray et al., 2012).Therefore glucose deficiency in the brain owing to altered vascular function is likely to be primary which results in altered plasma glucose dynamics mediated by autonomic inputs. Our model incorporates this possibility.

Impaired vascular function is known to be central to diabetic complications. The classical thinking has been that hyperglycemia causes the types of vasculopathies typical of T2DM. However, it has not been ruled out that vasculopathies are not primary. There is considerable evidence that microvascular alterations precede T2DM and are good predictors of it (Muris et al., 2012; Nguyen et al., 2007; Stehouwer, 2018; Zaleska-Żmijewska et al., 2017). Sedentary life style and deficiency of many specific types of physical activities and behaviors alter the expression of growth factors and endocrine mechanisms involved in angiogenesis (Watve 2013). It has been demonstrated repeatedly that many growth factors and angiogenic factors are responsive to behaviors, physical activity and exercise(Aloe et al., 1994; Cao et al., 2010; Chodari et al., 2016; Lakshmanan, 1986; Nexø et al., 1984; NEXØ et al., 1981; Tirassa et al., 2003). Deficiency of these behaviors is likely to lead to a primary deficiency of angiogenic mechanisms(Watve, 2013). We consider the possibility that altered vascular function in the brain is the primary reason why glucose transport from blood to brain is reduced. This creates chronic glucose deficiency in the brain to which the brain reacts by influencing multiple mechanisms of glucose regulation. The predominant mechanism is autonomic signaling. It is well known that sympathetic and parasympathetic tones are altered in T2DM(Thackeray et al., 2012). We consider the possibility that altered autonomic signaling is causal rather than consequential to diabetes, based on studies showing that changes in autonomic function precede T2DM (Carnethon et al., 2003).

The brain glucose uptake is insulin independent (Gray et al., 2014; Hasselbalch et al., 1999). Nevertheless insulin has many other functions in the brain and insulin signaling is known to be important in memory, cognition, decision making and behavior (Kerns 2001, Shemesh et al., 2012; Strachan, 2003). Brain is rich in insulin receptors and activation of certain cognitive functions in the brain is a likely cause of increased brain glucose utilization by insulin stimulation(Bingham et al., 2002; Rebelos et al., 2021). We assume therefore that brain has a fundamental requirement for insulin, independent of glucose metabolism and therefore mechanisms to ensure the required insulin levels in the brain are as much critical as ensuring minimum brain-glucose level. This is also ensured by autonomic mechanisms. Autonomic inputs are known to regulate β cell population and as well as insulin release from β cells (Thorens, 2015). Therefore, our model also incorporates brain control over insulin production.

Ensuring minimum supply of glucose as well as insulin to the brain is crucial during fasting when the plasma levels are low. The post meal levels of glucose and insulin in the plasma are much higher and at this time, the brain need not actively regulate the plasma levels of both. This can be taken care of by the peripheral mechanisms.

### 1.4 Objectives of the model

Our attempt is to construct a model whose predicted outcomes match with all or most of the observed phenomena listed above, qualitatively. We intend to construct a model in which different hypotheses for hyperglycemia can be used to make differential testable predictions. This approach can allow us to evaluate comparatively which causal factors individually or in combination can give us the set of predictions that match with the patterns listed above.

Out of the parameters required for the model, only some have empirical estimates available (table 1). In the absence of realistic estimates of all parameters, we do not intend to make a model making quantitative predictions. When a sufficiently large number of parameters can be optimized, it is not difficult to fit the data quantitatively. Therefore, we prefer qualitative predictions over quantitative ones. We test whether the model is able to predict the pattern observed empirically under some set of parameters. The ability to predict a given pattern is not a proof of the validity of the model, but the inability to predict an observed phenomenon at any set of parameters in a realistic range is a strong reason to call the model either inadequate or wrong. If an additional consideration compatible with the model is able to explain a pattern not explained by the main model, the model can be called inadequate but not falsified. However, if the model outcome directly contradicts a consistent and reproducible empirical finding, it can be considered falsifying evidence. We emphasize the need to evaluate our model in comparison with classical models on these lines.

**Table 1:**
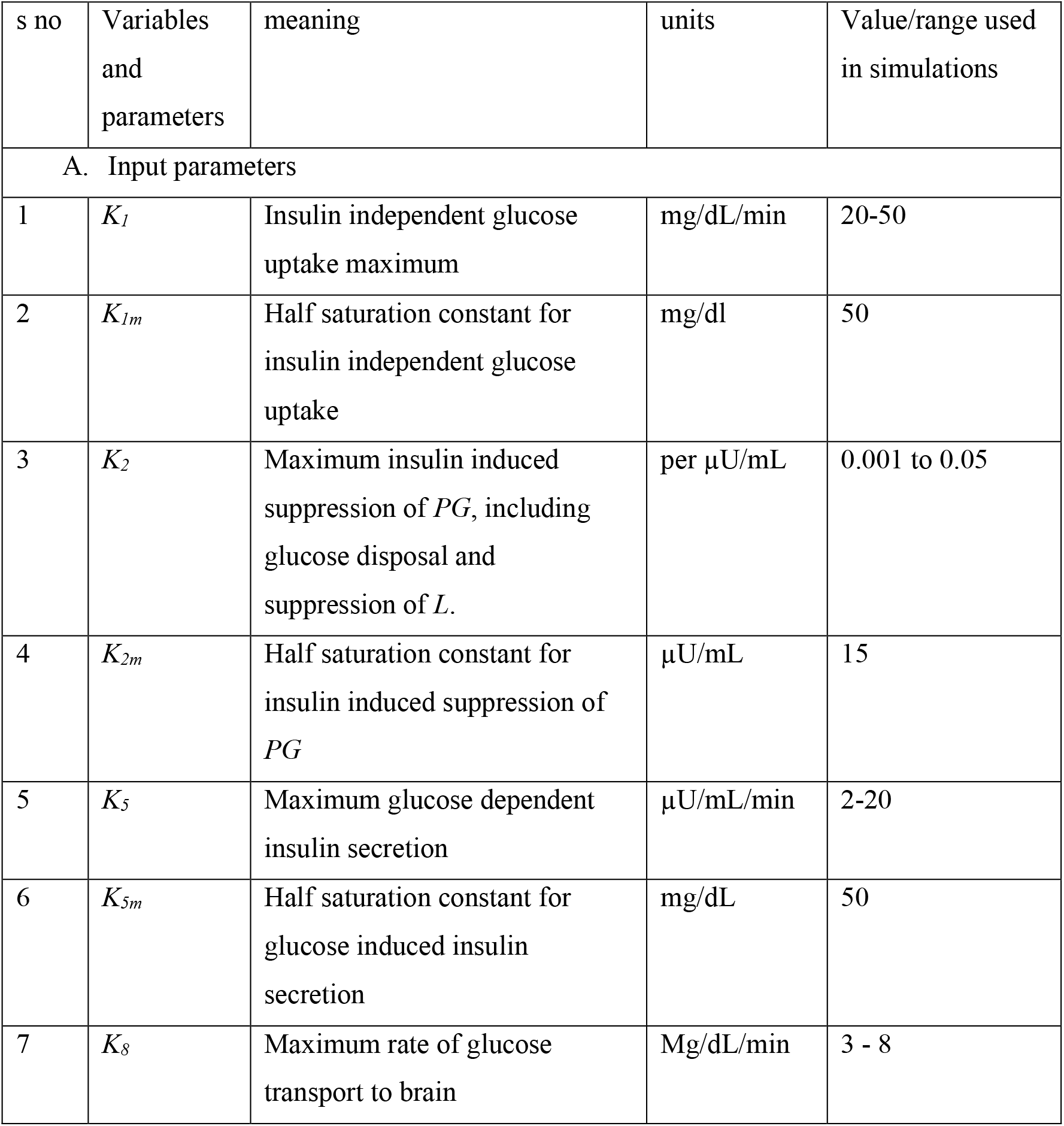
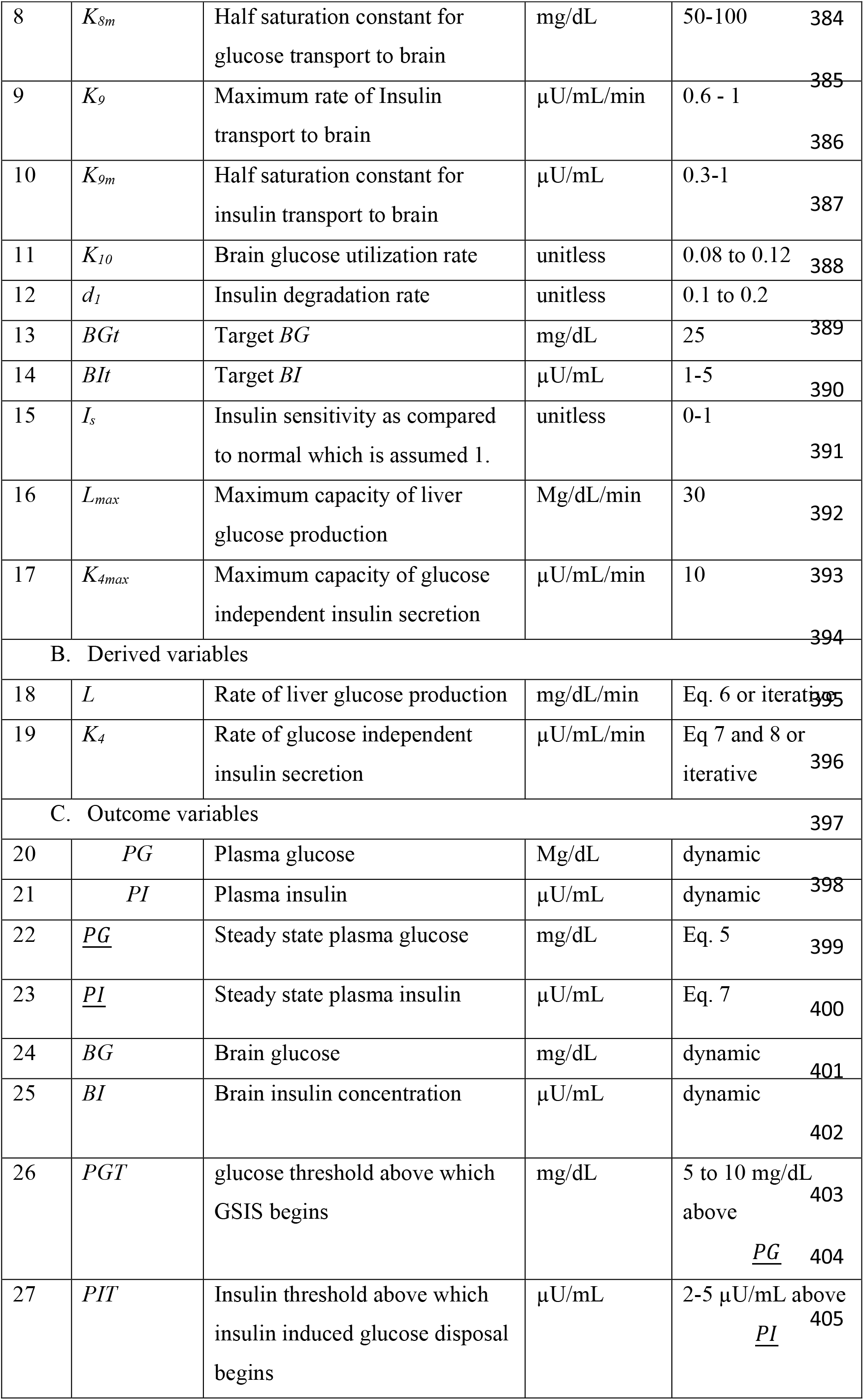
all variables and parameters considered in the model.

## 2. Methods

### 2.1 Assumptions of the model

A unique assumption of our model based on the analysis by Diwekar-Joshi and Watve(2020) is that the mechanism of regulation of glucose and insulin is different in the steady state and post meal state. In the steady state the central mechanisms are more important whereas in post meal state mainly the peripheral mechanisms are at work. Insulin induced glucose uptake and insulin dependent inhibition of liver glucose production happen only above a threshold glucose and insulin level respectively. The presence of such thresholds and their context dependent variability has been known for a long time (Chen et al., 1993; Henquin et al., 2006; Sorensen 1985). Below the threshold, other mechanisms regulate the levels (Sorensen 1985). We assume in the model that the thresholds are under neuro-endocrine control and fine-tuned by the brain. The thresholds found in isolated cell cultures are slightly lower than the normal fasting blood sugar and it is difficult to estimate thresholds in vivo (Chen et al., 1993; Henquin et al., 2006, 2015). In presence of autonomic signals, we assume the thresholds to be above the steady state target levels. In conditions under which the fasting levels need to increase, we assume the thresholds to increase proportionately. This may happen with peripheral or central mechanisms. These assumptions about the threshold make the model a TSS model.

The steady state glucose is decided by a balance between basal level of liver glucose production which is regulated by neuronal mechanisms and insulin independent glucose uptake by tissues. The steady state insulin level is decided by basal rate of glucose independent insulin production, which is regulated by neuronal mechanisms against the insulin degradation rate. Glucose is transported from plasma to brain by facilitated diffusion proportionate to capillary surface area in the brain and glut-1 expression. Steady state brain glucose is determined by glucose transport to brain and utilization by brain tissue. Brain needs a target level of glucose and the brain ensures it by directly regulating liver glucose production by neuronal mechanisms.

Insulin has cognitive and other functions in the brain(Begg & Woods, 2013; Kern et al., 2001; Rohner-Jeanrenaud & Jeanrenaud, 1983; Shemesh et al., 2012; M W J Strachan, 2005), for which the brain requires a target level of insulin, which it ensures by autonomic regulation of β cell number and basal insulin secretion independent of glucose. For ensuring the brain target level for insulin, a minimum plasma insulin level needs to be maintained. Maintenance of this level is assumed to be independent of glucose stimulated insulin secretion.

The rate of transport of glucose and insulin from blood to brain is not constant. Both are affected by the capillary density and blood flow. But they also depend upon specific transport mechanisms. Glucose is transported across the blood brain barrier mainly through the specific transporter glut-1. The expression of glut-1 in brain and other tissues is variable and dependent on multiple dietary, endocrine and growth factor related mechanisms (Boado et al., 1994; Ge et al., 2011; Liu et al., 2018; Schüler et al., 2018).Glucose transport from blood to brain is diminished in obesity and prediabetes (Hwang et al., 2017). Insulin transport to brain is also reduced in obesity (Begg, 2015). The ratio of CSF to plasma insulin is inversely proportionate to obesity and insulin resistance(Gray & Barrett, 2018). More over inducing endothelial dysfunction and reducing glucose transport experimentally by endothelial specific deletion of HIF 1-α resulted into hyperglycemia (Huang et al 2012). Based on such multiple lines of evidence we incorporate the possibility of diminished glucose and insulin transport to brain being the primary pathology of T2DM and hyperglycemia only a consequence.

#### The model

Since we assume that the steady state and post meal mechanisms of glucose regulations are not identical, we model and analyze the two states in two phases of the model. The variables and parameters used in the model and the range of parameter used are in table 1.

##### A: Modeling steady state glucose

As per the assumption, below a threshold *PI*, insulin induced glucose uptake and insulin induced inhibition of liver glucose production is negligible. Therefore, we write,

1. Plasma Glucose:

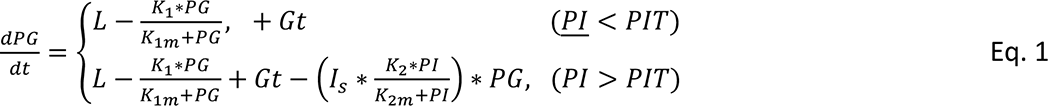
2. Plasma Insulin:

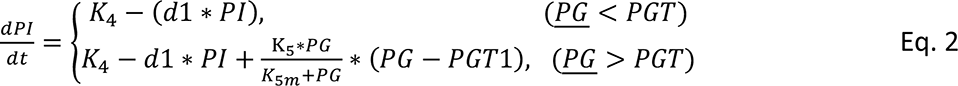
3. Brain glucose:

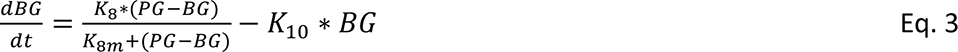

Since brain forms only about 3 % of total body mass, we assume plasma glucose diffused to the brain is a negligible fraction of total plasma glucose. Therefore, that term is not included in the plasma glucose dynamics. Similarly, insulin diffusion to brain is not represented in the plasma insulin dynamics. Nevertheless, the rate of plasma to brain diffusion is crucial in determining the brain glucose and insulin dynamics.

4 Brain insulin:

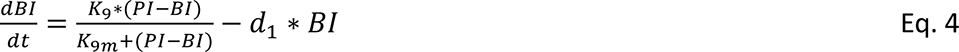

At steady state (SS) all differential terms become zero and steady state levels can be calculated using equilibrium solutions.

> As per the assumption of the model the brain needs a certain level of glucose and that is the target of homeostasis *BGt.* In order to ensure *BGt* at given K_8_, K_10_ and K_8m_, the SS plasma glucose *<u>PG</u>* needs to be

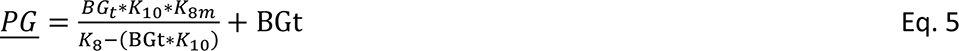

> We assume that the brain ensures this plasma glucose level by regulating liver glucose production *L*. To maintain the desired plasma glucose level at steady state, the liver glucose production should be:

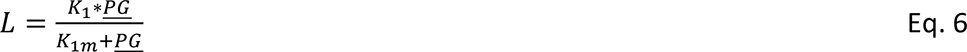

> The brain can either ensure this by sending a calculated signal to the liver through the autonomic system. Alternatively, this can be ensured by the brain by increasing sympathetic inputs when *BG <BGt* and parasympathetic when it is above target(Antuna-Puente et al., 2009; D’Alessio et al., 2001; Kiba, 2004). The required *L* can be achieved iteratively by 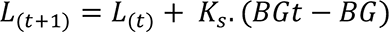

> Similarly, to ensure required brain insulin levels, plasma insulin should be

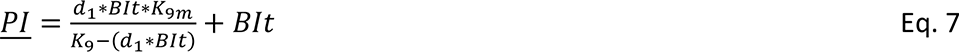

> To ensure this much plasma insulin level, K4 should be

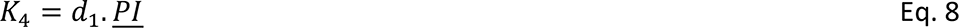

> Similar to glucose, the brain can ensure the target level of insulin in the brain by neuronally modulating *K_4_* in a calculated or iterative manner. Specific autonomic inputs are known to regulate β cell number as well as insulin release from β cells(Begg& Woods, 2013; Boland et al., 2017; Kiba, 2004).

We also assume that both L and K_4_ have a fixed upper limit as L_max_ andK_4max_ when the maximum capacity of liver and pancreas is reached. L and K_4_ cannot exceed this limit even if neuronal inputs become more intense. This happens when *K_8_* declines below a threshold such that,

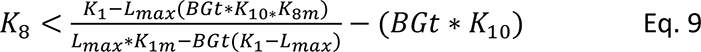

There is a potential conflict here. Sympathetic tone is known to increase *L*(Nelles et al., 1996; Nonogaki, 2000)whereas parasympathetic signal is required for proliferation of β cells. Simultaneously sympathetic signal is known to inhibit insulin release from β cells(Gilon & Henquin, 2001; Miller et al., 1976). If the brain is deficient in glucose as well as insulin, both sympathetic and parasympathetic tones will be simultaneously higher(Watve et al., 2014). This is incorporated into the model by updating 𝐾3 iteratively according to both *BG* and *BI* levels simultaneously,

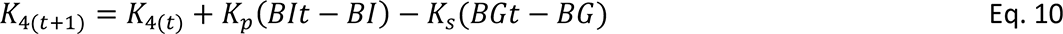

By this consideration if sympathetic stimulation of liver glucose production is adequate to restore required *BG*, 𝐵𝐺𝑡 − 𝐵𝐺 will tend to zero and there will be no interference in insulin release by β cells. However, in case the rise in liver glucose production is inadequate to ensure required *BG*, 𝐾3 will be affected which will apparently influence β cell function.

##### B. Simulating glucose tolerance curve

While for the steady state, equilibrium solutions can be algebraically derived, the post glucose load curve and its properties can only be obtained by simulations. Simulations were run using the same set of equations and giving a positive ephemeral *Gt* to simulate food intake.

In all the simulations used, insulin resistance can be simulated by altering *Is*, β cell dysfunction by reducing *K_4_* and/or *K_5_* and vascular defect slowing down transport of glucose and insulin to brain and other organs by altering *K_8_*, *K_9_* and *K_1_*. Mental stress is assumed to increase *K_10_*. By differentially altering these parameters the model can separately and collectively examine the effects of insulin resistance, β cell dysfunction, reduced rates of diffusion across BBB and stress on the glucose tolerance curve.

###### T2DM-causal analysis

We ask the question which minimal set of changes can give rise to stable increase in fasting glucose, changes in the glucose tolerance curve characteristic of type 2 diabetes along with other patterns listed above. The putative causal factors are examined individually and in combination. The factors include insulin resistance (Decreased *I_s_*), subnormal β cell response (Reduced *K_4_ and/or K_5_*), reduced insulin independent glucose uptake (decrease in *K_1_*), reduced blood to brain transmission of glucose (*K_8_*) and insulin (*K_9_*) and stress related increase in brain glucose consumption (*K_10_*). Subnormal or defective vasculature is expected to decrease *K_1_, K_8_* and *K_9_* proportionately, however the decrease in the three parameters may not be proportionate if the vasculature in different parts of the body is differentially affected. Also glucose transporters and there expressions can alter in tissue specific manner. Therefore we allow proportionate as well as differential decrease in the three parameters.

## Results

### A: Steady state solutions

We observed that the steady state solutions and the autonomic iteration simulations match well in the end result except when the limits of *L* and *K_4_* are reached. However, it took a long time to reach the desired level by iterative approach, particularly when the desired level was substantially different from the starting level. In real life major changes in vascular function or glucose transporter levels reflected in *K_8_*, *K_9_*, *K_1_* happen gradually. Therefore the desired level can be attained by autonomic fine tuning over time. Since in the iterative simulations, a long time was required to reach SS, we used the steady state solutions for quicker results.

It can be seen that the steady state levels of *PG* in the model is a stable steady state since substituting *PG <<u>PG</u>* leads to a positive value and *PG ><u>PG</u>* leads to a negative value of 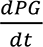 The same applies to plasma insulin, brain glucose and brain insulin. This ensures a stable steady state of glucose-insulin in the plasma as well as brain on fasting [1,2]. From equations 5, 6, 7 and 8 we see that, *FG* does not change by altering *Is* or *FI*. This is compatible with the empirical findings that insulin receptor knockouts and insulin suppression experiments fail to increase fasting plasma glucose [9]. This result is unique to our model and is due to segregating fasting and post meal glucose regulation mechanismsby using thresholds. This is the only explanation offered so far, for patterns [9], [16] and [17].

The steady state solution of our model shows that *FG* can increase as a result of increase in *K_10_* or decrease in *K_8_*. That is if the glucose consumption in brain increases and/or the rate of glucose transport from blood to brain decreases. Both the effects are interdependent and the shape of the glucose response is decided by the interaction between *K_8_* and *K_10_*. Increased *K_10_* increases *FG* marginally and almost linearly when *K_8_* is large. When *K_8_* decreases even a marginal rise in *K_10_* can induce disproportionately greater rise in *FG* (fig 1). This means that stress induced hyperglycemia is unlikely to be seen in healthy individuals while it is more likely in individuals with reduced vascular transport [7].

**Figure 1:**
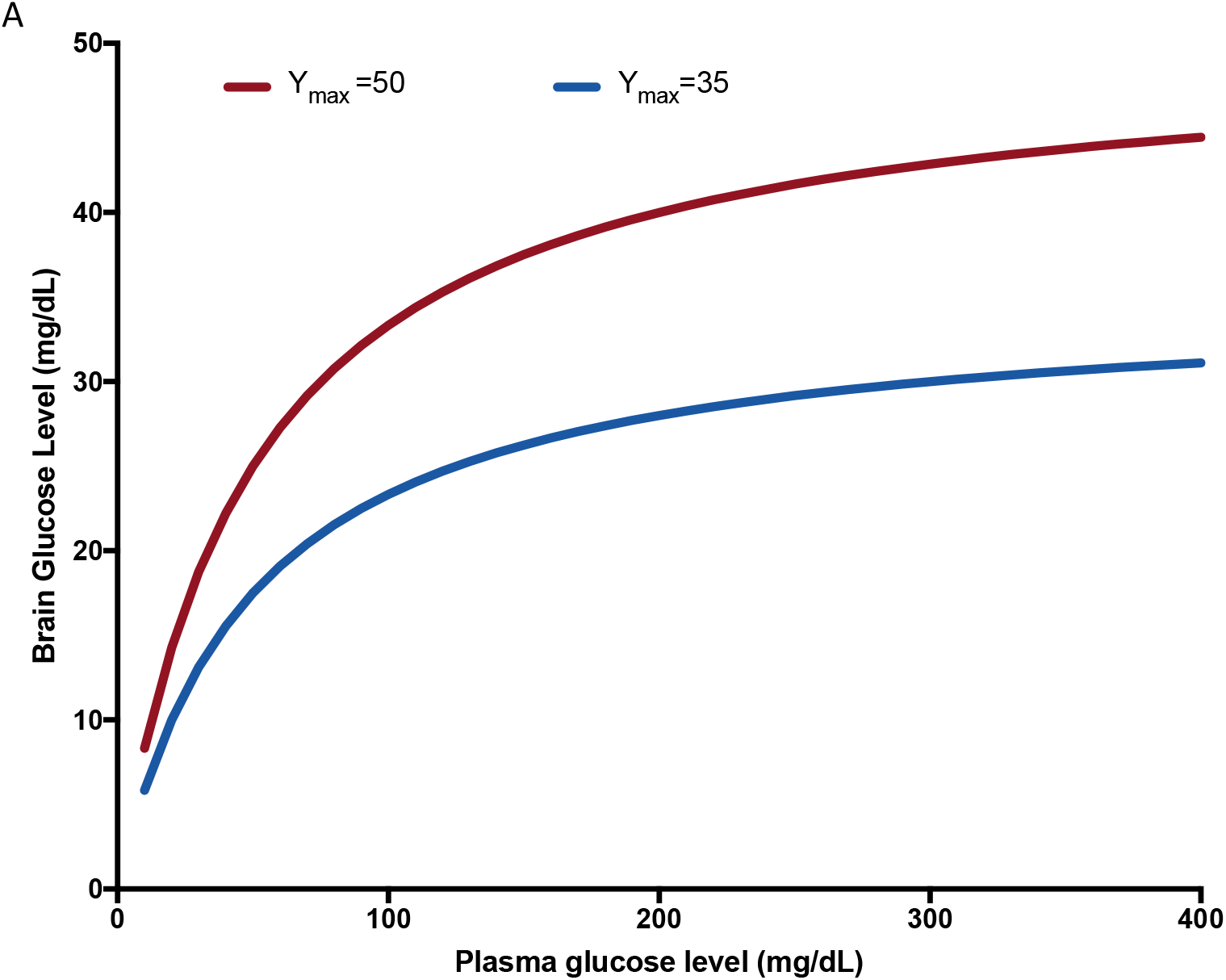
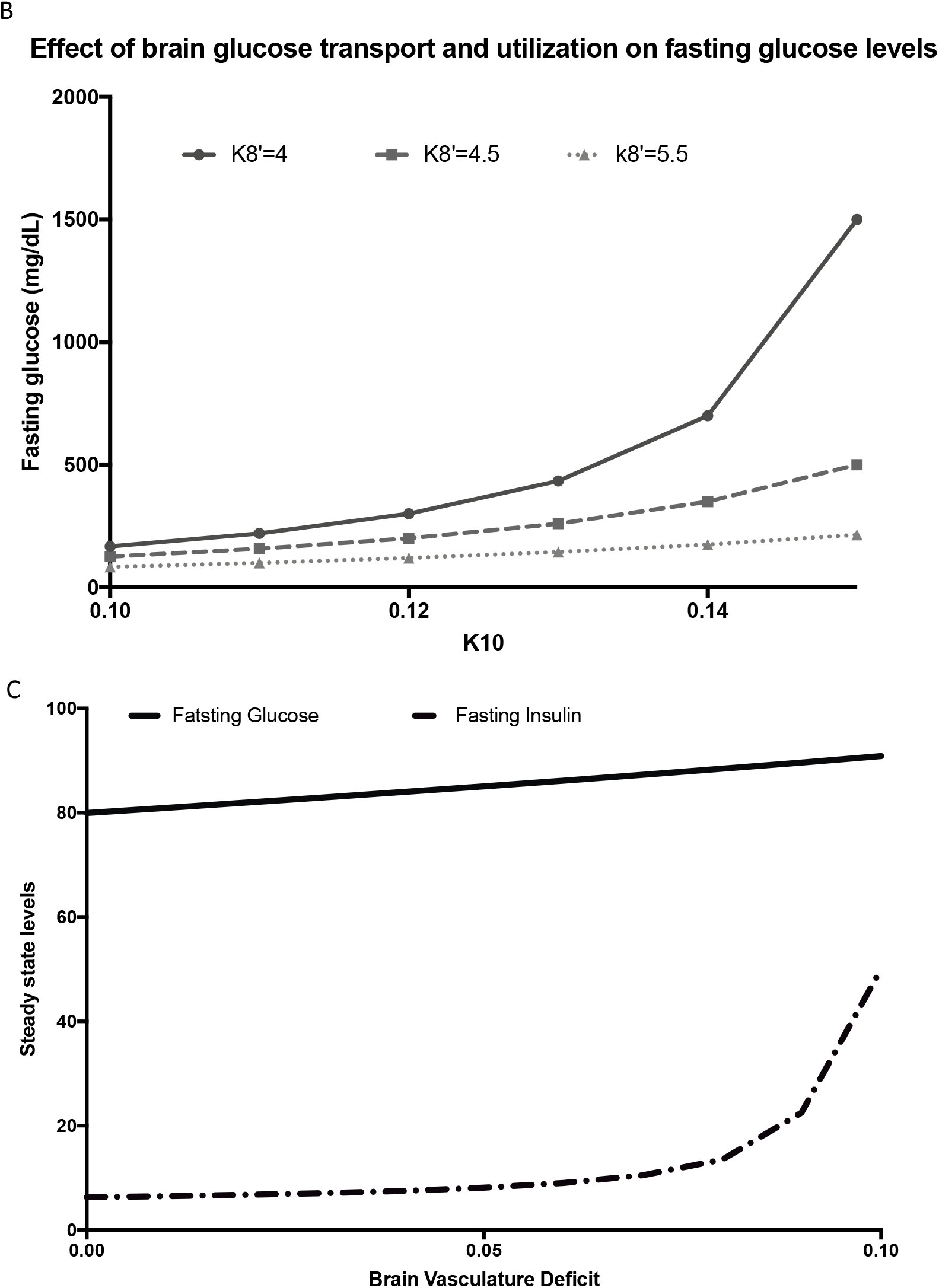
The same input different output phenomenon: A. A conceptual diagram with arbitrary units: A saturation relationship in glucose and insulin transport described by a Michaelis Menten type of equation has important consequences that can account for many phenomena observed in prediabetic and diabetic stages. For example, here for achieving an increase in Y by 6 units with a higher Y_max_, X needs to increase only by 6 units but at a lower Y_max_ for the same target increase in Y 25 units of increment in X is needed. This non-linearity explains many phenomena including prediabetic hyperinsulinemia and stress induced diabetes in our model of glucose homeostasis. B. A simulation using the steady state model: It can be seen that as K_8_ decreases, for the same change in brain requirement K_10_, a non-linear escalated increase in plasma glucose is required. C. A simulation result assuming a correlated decrease in K_8_ and K_9_. It is possible that while glucose shows a marginal increase, insulin increases substantially, since the parameters of glucose and insulin transport curves are different. This is a potential explanation for hyperinsulinemic normoglycemic state.

The origin of hyperinsulinemic normoglycemic condition can be explained by the difference in the parameters of the saturation curves of transport dynamics of glucose and insulin. If the insulin diffusion is assumed to be nearer to saturation and glucose diffusion is sufficiently away from saturation, at the same level of vascular function deficiency, insulin will increase disproportionately more than glucose (Figure 1). This is a possible cause of hyperinsulinemic normoglycemic state [3], and the reason why hyperinsulinemia precedes hyperglycemia [4] which requires neither insulin resistance nor compensatory hyperinsulinemic response. When this happens, both HOMA-IR and HOMA β increase although there is no change in true insulin resistance and compensatory insulin response. This gives a false impression of insulin resistance as clinically defined.

Further as the transport saturation constant *K_8_* and *K_9_* continue to decrease proportionately, *FG* increases monotonically but *FI* shows a non-monotonic curve in which *FI* increases with moderate decrease in transport rates but decreases after a threshold decrease in transport rates. This non-monotonicity is predicted by the iterative autonomic inputs model. When the required *L* approaches or exceeds *L_max_*, the steady state *BG* remains smaller than *BGt*. The resultant increase in the sympathetic signal inhibits insulin release from β cells. This leads to reduced insulin secretion and insulin levels drop substantially. Thus, without bringing in additional factors the model explains the early phase rise in fasting insulin as well as the later phase decline in the course of T2DM [4] (Figure 2).

**Figure 2:**
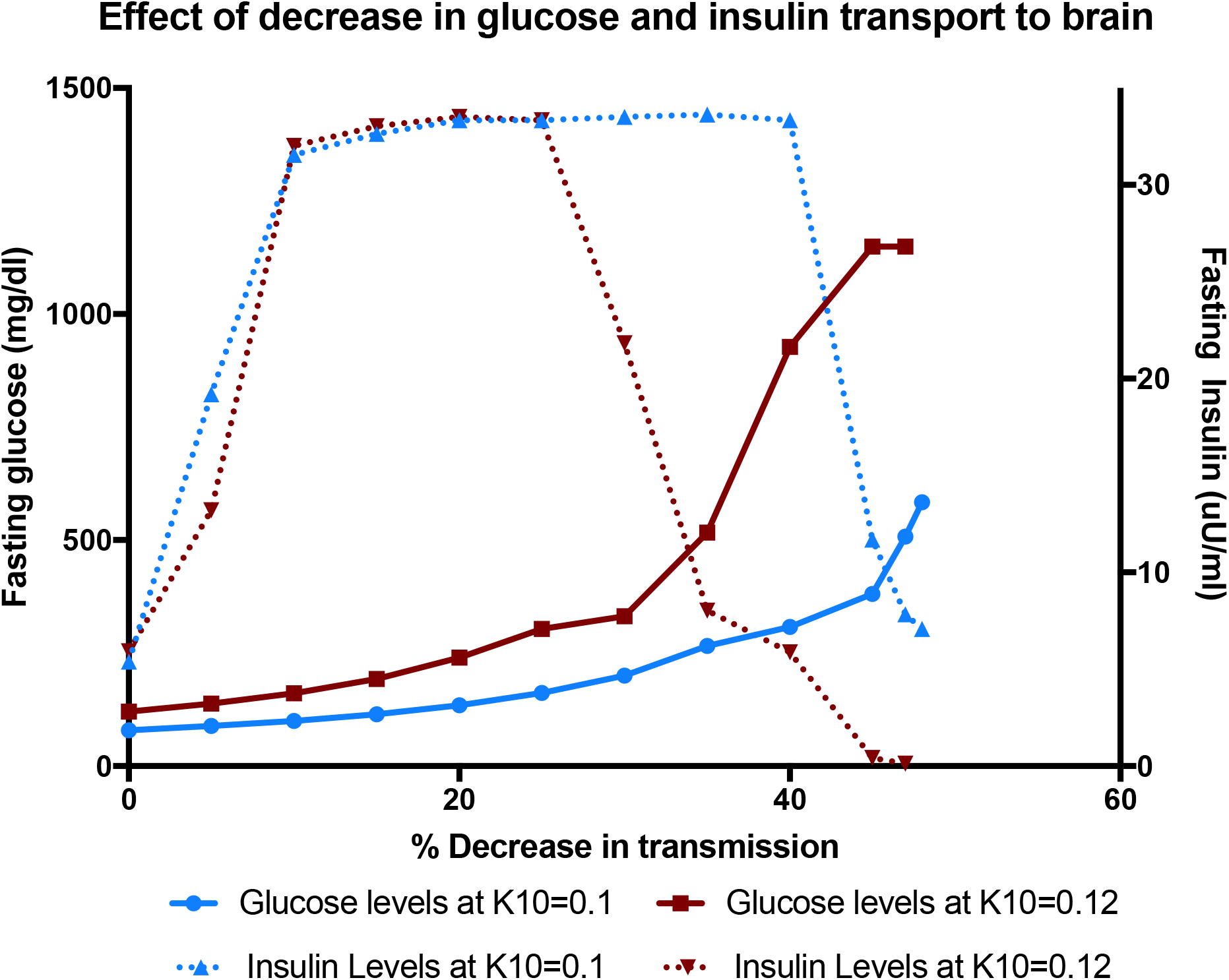
Effect of correlated decrease in K_8_ and K_9_ on fasting plasma glucose and insulin. Note that for a small to moderate decrease FI increases rapidly with marginal change in FG. However, after a threshold decrease in the transport rates FI declines sharply whereas FG increases rapidly. This pattern matches with the course of clinical T2DM without involving insulin resistance, compensatory hyperinsulinemia and β cell exhaustion. Simulation specific other parameters were K_1m_ =50, K_1_=30, K_2m_=15, K_2_=0.01, K_5m_=0.01, K_5_=0.5, d_1_=0.15, I_s_=1, BGt=25, BIt=5, K_8m_=80, K_8_=5, K_9m_=0.3, K_9_=0.8, K_10_=0.1(Blue line) and 0.12(Red line).

This decline in the insulin response does not require β cell dysfunction as a causal mechanism but there are more complex possible effects on β cells. Simultaneous activation of sympathetic and parasympathetic inputs to β cells is implicated in β cell amyloidogenesis (Watve et al., 2014). Parasympathetic stimulation is known to increase β cell number. However, sympathetic signaling suppresses insulin release resulting in increased retention time of insulin along with amylin which beyond a threshold retardation may result in to spontaneous polymerization of amyloid protein leading to poisoning of β cells and amyloidogenesis (Watve et al., 2014). This is an alternative causal interpretation of β cell dysfunction and decline in its population. This mechanism has a built-in negative feedback loop giving rise to a steady state β cell population. This result is compatible with the finding in which the β cell population remains subnormal lifelong [11,12]. This is in contrast with the dynamics expected by the classical thinking in which β cells are destroyed by gluco-lipotoxicity or oxidative stress. This mechanism has a built-in positive feedback cycle. As a part of the β cell population is destroyed, the insulin secretion is decreased which would increase glucose further accelerating gluco-lipotoxicity and oxidative stress and therebyβ cell loss. Such a positive feedback vicious cycle can only stop with complete destruction of the β cell population. This prediction of the classical model does not match the finding of sustained presence of subnormal β cell population in T2DM.

By the classical model, prolonged sustained exercise in the fasting state should result into lower plasma glucose. However, experimental increase in glucose uptake resulting from physical activity during fasting does not necessarily result into a decrease in *FG*. This is mainly because as *FG* starts decreasing, *BG* decreases consequently. *BG* being lower than *BGt* induces sympathetic mediated increase in *L* and the change in *FG* by physical activity is more or less compensated. This may happen through the agency of glucagon as well, since a strong link between autonomic signals and glucagon secretion is known. Therefore, at times plasma glucose may actually increase after intensive exercise. Further, if exercise involves heightened activation of cerebellar mechanisms of coordination, *K_10_* may also increase resulting into higher rather than lower *FG* in response to exercise [8].

A high dose of exogenous insulin results into increased plasma glucose after a time lag, is known as Somogyi phenomenon. The phenomenon is conditional and not seen in every diabetic. It is said to involve the counter-regulatory response on facing central hypoglycorrhachia. However, often this response is seen without peripheral hypoglycemia. This is best explained by reduced *K_8_* that gives rise to decreases *BG/FG* ratio. With smaller *K_8_,* at apparently normal or increased plasma glucose, brain glucose can still be lower than the target which gives a sympathetic signal to increase *L*. Simulations using the autonomic iteration model shows this phenomenon quite well at low *K_8_*. At the same level of insulin infusion, at lower *K_8_* there is more intense hyperglycemic response (Figure 3) [14]. A model without accommodating changes in *K_8_* does not predict Somogyi phenomenon without an obvious hypoglycemic state.

**Figure 3:**
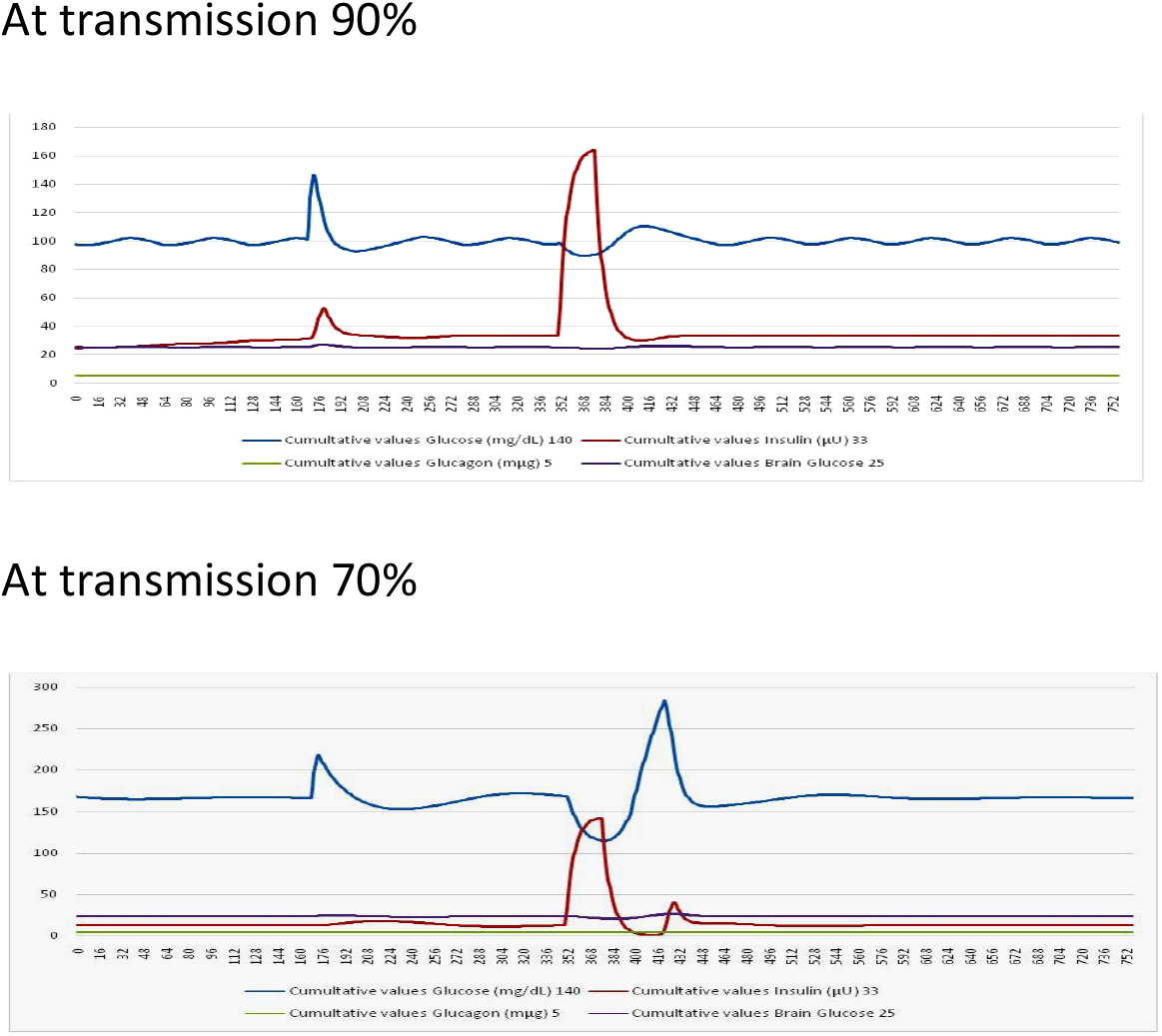
Effect of high dose insulin on peripheral glucose under two rates of glucose and insulin transport across the blood brain barrier (Correlated decrease in K_8_ and K_9_). A paradoxical rise in blood sugar following insulin administration is more prominent when the transport rate is substantially lower than normal. Other parameters (K_1m_ =50, K_1_=30, K_2m_=15, K_2_=0.08, K_5m_=0.01, K_5_=15, d_1_=0.15, I_s_=1, BGt=25, BIt=4.5, K_8m_=80, K_8_=5, K_9m_=0.3, K_9_=0.8, K_10_=0.1)

The response is restricted to a narrow set of conditions. For intense hyperglycemic response to insulin administration, it is necessary that insulin sensitivity is good, *BG* is close to *BGt* before insulin transfusion, and *K_8_* is subnormal.

Potentially one can visualize two distinct possible classes of reasons why blood sugar increases in response to brain injury. In the first, there is some impairment in the glucose regulation mechanism. In the second, the requirement for glucose is increased during the wound healing process and hyperglycemia is a mechanism of the body to meet the demand. It is known that infections such as meningitis, stroke or hemorrhage leads to transient hyperglycemia. Such hyperglycemia is often accompanied by lower *BG* levels (Schut et al., 2009; van Veen et al., 2016)[15]. In such cases, hypoglycemic treatments can be counterproductive and result into delayed or derailed repair process. This is a likely reason why in patients under intensive care, strict regulation of plasma glucose increased mortality instead of the expected decrease (The NICE-SUGAR Study Investigators, 2009).

When hyperglycemia is a process of meeting increased glucose demand or compensating subnormal transport, removal of plasma glucose by any means is expected to increase *L*. This is observed to happen when SGLT2 inhibitors decrease the urinary threshold and drive out plasma glucose. If increasing *L* is not sufficient to restore the required glucose level, there is a shift to ketogenesis since the brain can use keto acids as alternative source of energy. Therefore, increased *L* with or without increased ketogenesis is expected following SGLT2 inhibitors [13].

### B: Glucose tolerance curve

Unlike the steady state predictions, the patterns of post feeding glucose curve by our model are not qualitatively different from classical models. The area under the curve, height of the peak, time required to return to steady state and the time difference between glucose and insulin peaks are increased by decreased *I_s_*, *K_5_*, or *K_8_* individually or in combination. Decrease in *I_s_* or *K_5_* does not increase *FG* but changes the shape of the glucose tolerance curve. Decreased *K_8_* may alter both simultaneously. The altered curve shows the three typical features namely taller peak, delayed return to steady state and longer gap between glucose peak and insulin peak [5]. This result is not unique to our model and classical models also show the three features. The classical model results into simultaneous and proportionate alterations in the fasting as well as post meal glucose and therefore fails to explain the NFG-IGT state. In our model, decreasing *I_s_* or *K_1_* without a change in *K_8_* results into an NFG-IGT state [6].

### C: Population simulations

For the steady state as well as post glucose load dynamics we give population distributions to *K_1_, K_5_, K_8_, K_9_, I_S_* and also incorporate normally distributed error in glucose and insulin measurements in fasting and post meal sampling. We also incorporate correlated changes in *K_1_, K_8_* and *K_9_* which are expected as a result of hypo-vascularization in the brain. These simulations are run to observe whether we get the anomalous correlations in fasting versus post meal state and in the HOMA indices as observed empirically (Chawla et al., 2018; Diwekar-Joshi & Watve, 2020).

By classical models the regression correlation parameters of glucose-insulin relationship are not different in fasting state versus post glucose load although the range of variables is different as shown previously by Diwekar-Joshi and Watve (2020). Also, if we assume HOMA-IR to faithfully reflect insulin resistance and HOMA β to faithfully represent β cell response, then there is no reason why the two indices should be correlated.

The assumption behind our model that there are different mechanisms at work under fasting versus post glucose load condition is necessary to explain the large difference in the regression correlation parameters in fasting versus post meal levels. If the same set of mechanisms in the fasting and post meal conditions are operational, whatever the model used, it is imperative that fasting correlation regression parameters are stronger or comparable to post meal parameters. Simulations with our model are able to give poor *FG-FI* correlation along with strong post meal glucose insulin correlation under multiple conditions [16] (Figure 4). When the variance in *K_8_* and *K_9_* is small but that in *K_1_*, *K_5_*, *Gt* and *I_s_* individually or in combination is large, the fasting correlations are weak and post meal correlations strong; the post meal glucose-insulin regression slope is substantially greater than *FG-FI* slope. Also, whenever *FG-FI* correlation is weak, the indices HOMA-IR and HOMA β are strongly correlated similar to the epidemiological data [17]. The difference between fasting and post meal regression correlation patterns is not predicted by the classical models and is unique to our model which assumes different set of regulatory mechanisms in the fasting and post meal state.

**Figure 4:**
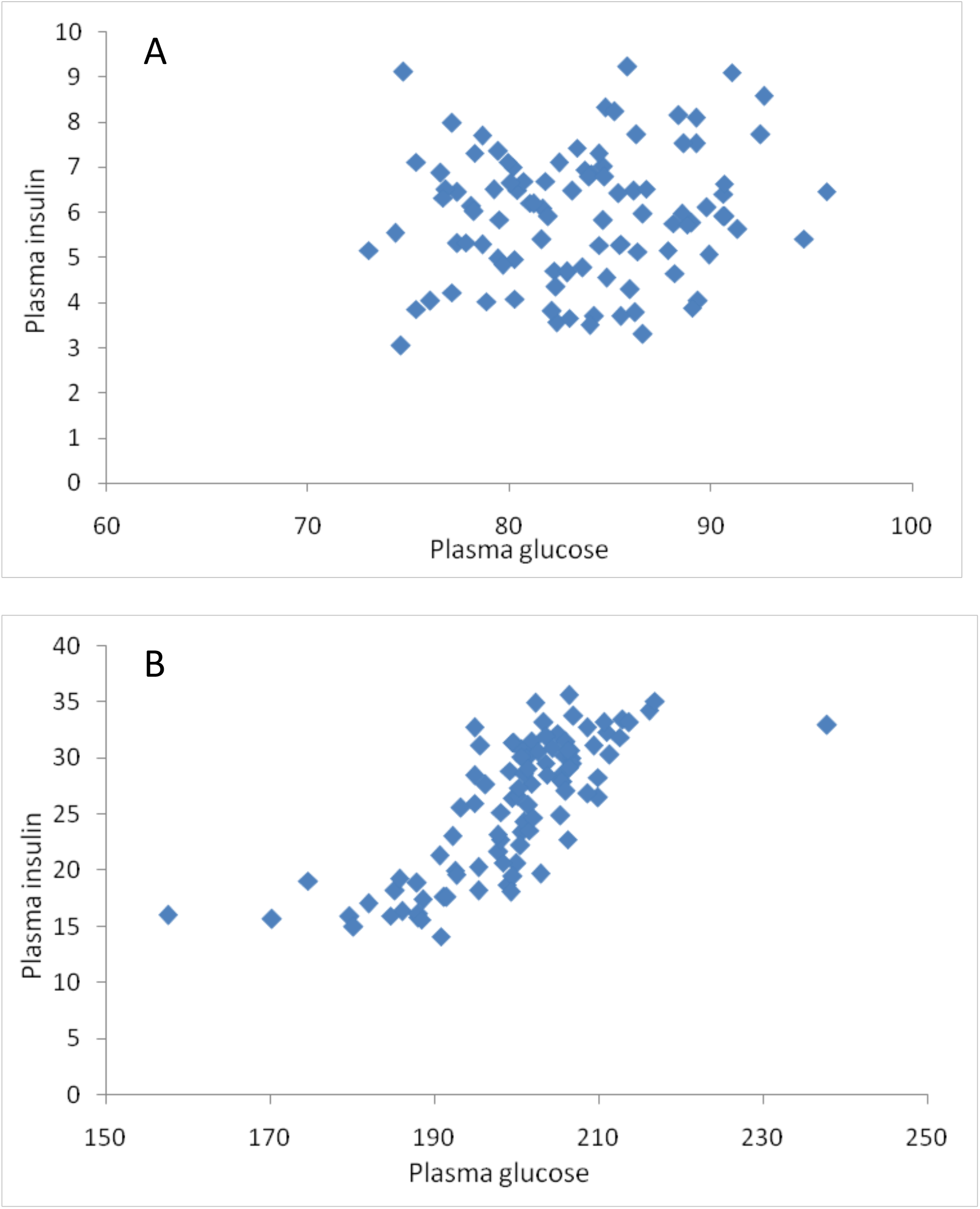
Plasma glucose and insulin correlation in the fasting steady state (A) and post meal (B) in simulations. The empirical finding that there is a strong positive correlation in the post meal data but poor correlation in fasting steady state was possible under a variety of conditions in our model. Depicted here the simulation results (post meal R^2^=0.57, steady state R^2^=0.014) in which only K_1_ was given a wider population distribution. (mean (S.D.) K_1_ = 35(7), K_5_ = 10(0.00001), K_8_ = 5(0.000005), and K_9_ = 0.8(0.000008). Other parameters (K_1m_ =50, K_2m_=15, K_2_=0.05, d_1_=0.15, I_s_=1, BGt=25, BI=4.5, K_8m_=80, K_10_=0.1).

### D: Effect of glucose normalization on arresting diabetic complications and mortality

By classical thinking, chronic hyperglycemia is responsible for the diabetic complications and preventing hyperglycemia should arrest complications. In contrast the thought behind our model is that vascular problems are primary which alter the rate of glucose insulin transport to the brain and hyperglycemia is an offshoot symptom that may not be causal to diabetic complications. The diabetic complications can arise directly from the vasculopathy. Therefore, controlling glucose may not have any effect on diabetic complications [18]. On the other hand, forcefully reducing plasma sugar without addressing vascular problems can create more severe glucose deficiency in the brain and other organs, thereby turning counterproductive. Because of the saturating dynamics of transport, a curvilinear relationship is expected between plasma glucose and brain glucose in such a way that moderate reduction in hyperglycemia will change brain glucose availability marginally whereas tight glucose regulation can have disproportionately larger effect (Figure5). Therefore, tight glucose regulation may increase mortality and other adverse outcomes. This prediction is compatible with some of the tight versus moderate control clinical trials including ACCORD, NICE sugar trial and UGDP [18].

## Discussion

The main inference from our model, stated most conservatively, is that a brain centered model can potentially explain most of the anomalies faced by peripheral models and therefore needs greater attention (Figure 6). If supported well, by exploring its testable predictions suggested below and possibly more, it has a potential to bring in a radical change in the fundamental view as well as clinical practice of T2DM.

**Figure 5:**
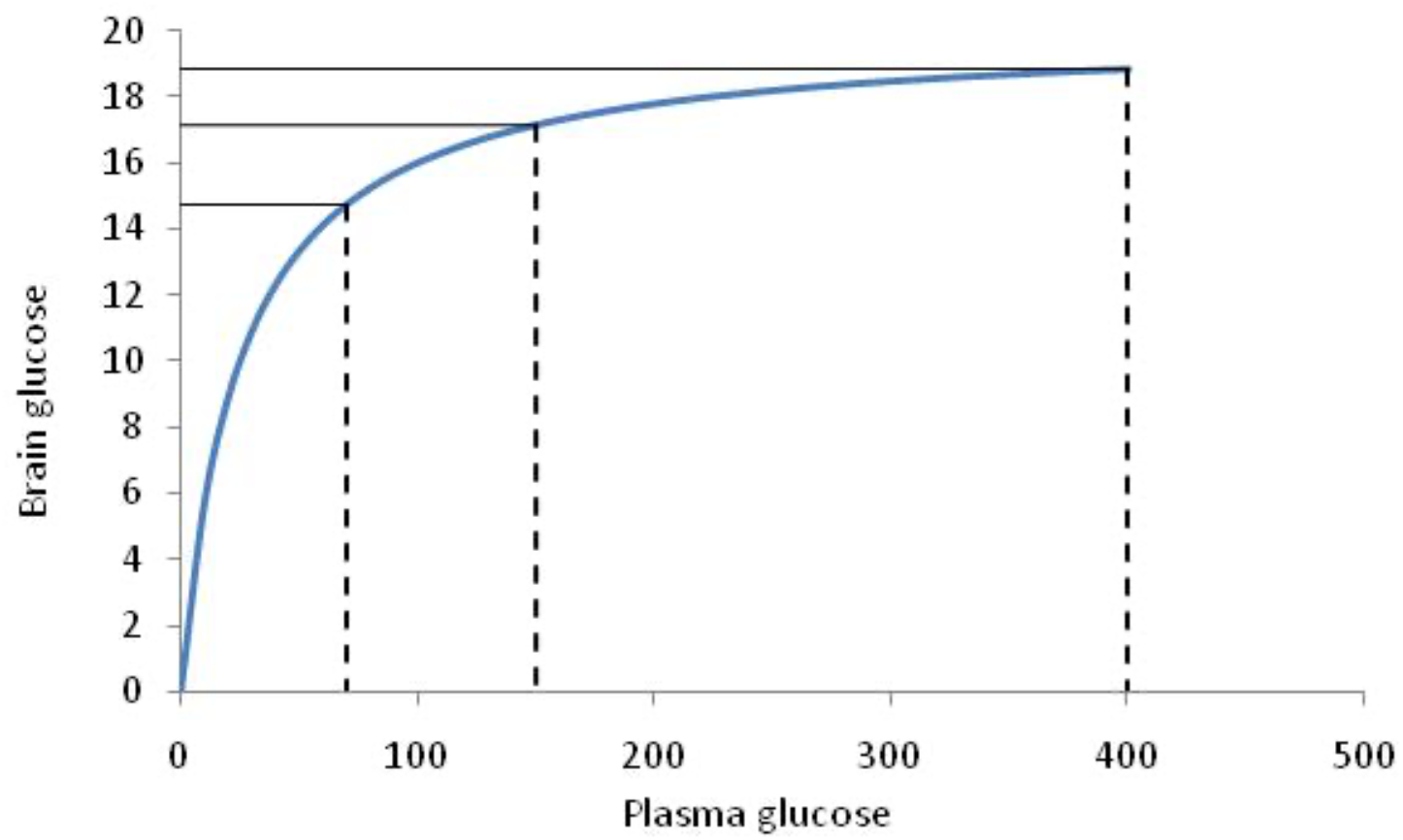
Effect of glucose lowering on brain glucose availability. For example, a reduction in FG from 400 mg/dl to 150 mg/dl corresponds to a decrease in BG by1.68 mg/dl, but further reduction from 150 to 70 mg/dl reduces BG by 2.41 mg/dl. Therefore a moderate sugar control may not affect brain glucose supply drastically but tight sugar control is expected to affect it more seriously.

**Figure 6:**
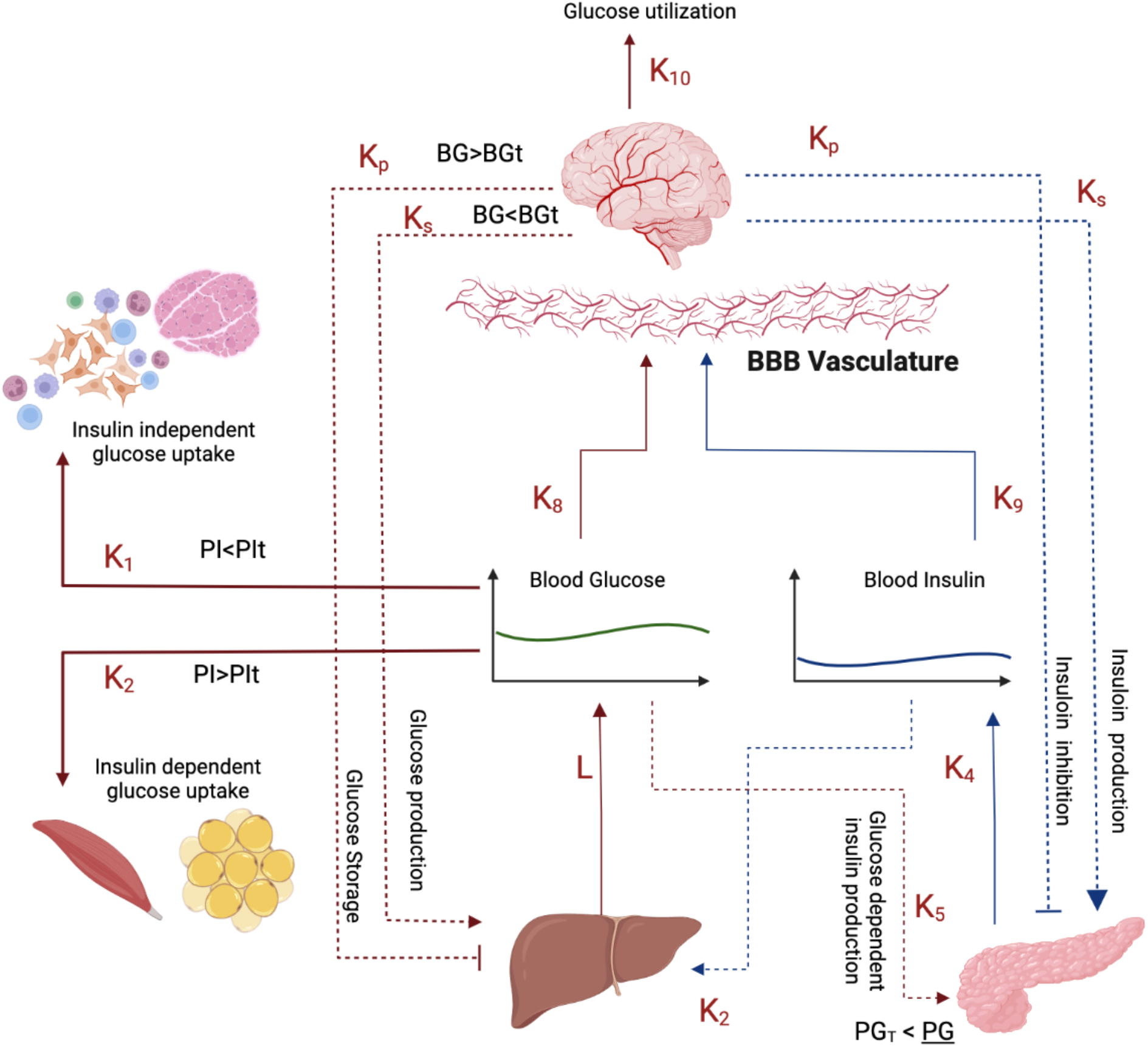
Effect of changes in vasculature in blood brain barrier. The model predicts that glucose and insulin transportation to the brain has proportionately major role in development of diabetic symptoms compared to other peripheral changes. The blood brain barrier vasculature hence may hold the key to understanding the shift from normal to diabetic condition. (created using BioRender)

### Testability of assumptions and additional predictions

The assumption that the thresholds *PGT* and *PIT* can be modified by autonomous signaling needs to be tested empirically. Although currently the thresholds are known to be flexible, information about the conditions and mechanisms of change are poorly known. Our assumption that the parameters of saturation equation for glucose and insulin transport to the brain are different and under normal conditions insulin transport is closure to saturation than glucose transport can be tested with carefully worked out kinetics of the two transport mechanisms.

The assumption of our model that reduced transport of glucose and insulin to brain is the primary pathology of T2DM leading secondarily to hyperglycemia makes more predictions that can be tested experimentally or epidemiologically. Experimentally specifically blocking glut1 receptors in the brain should lead to hyperglycemia. Conversely infusion of glucose directly to the brain should reduce peripheral hyperglycemia in the short run. This is already suggested by some experiments (Ono et al., 1983; Osundiji et al., 2012). It is also demonstrated that inducing primary endothelial dysfunction and reduced glucose transport to brain by endothelial deletion of hypoxia inducible factor HIF-1α results in hyperglycemia (Huang et al., 2012). More careful research in this direction to reveal the cause effect relationship between vascular defects, brain glucose levels and plasma glucose levels will be enlightening. Epidemiologically hypoglycemia is shown to associate with dementia and other neuronal problems (Lipska & Montori, 2013; Meneilly & Tessier, 2016; Rhee, 2017; Yaffe, 2013), tight glycemic control led to higher mortality as compared to moderate control in many of the clinical trials (Klein, 2010; Schwartz & Meinert, 2004; The NICE-SUGAR Study Investigators, 2009) which demands investigations into the causal pathways. The question whether tight glycemic control leads to subtle neuronal changes in the long run as expected by our model needs careful investigation. The difference between fasting and post meal regression correlation parameters between glucose and insulin is an important epidemiological line of evidence we have used. Chawla et al (2017) and Diwekar-Joshi and Watve (2020) showed this pattern across four different data sets. How generalized the pattern is needs to be tested in multiple population data.

On the modeling front it is necessary to undertake comparative evaluation of the different models with respect to the battery of prediction that we listed here. Perhaps a few more predictions may be added. However, at present many of the models and their possible modifications are not explored sufficiently to see whether they can explain the currently unexplained patterns under certain set of conditions. A model prediction matrix would be an appropriate approach for such a comparative evaluation, but we may have to wait till all alternative models are explored sufficiently elaborately on which of the empirical patterns they predict, which ones they are compatible with and which ones they contradict. We have shown here that the brain centered model predicts or is compatible with all the patterns listed in the introduction and does not contradict any.

### Possible causes of T2DM

The classically believed causal factors namely insulin resistance and β cell dysfunction are not compatible with many of the empirical findings as shown by Diwekar-Joshi and Watve (2020). In our model change in insulin resistance and β cell dysfunction were neither necessary nor sufficient to account for all the patterns. Nevertheless, they were helpful in accounting for the altered glucose tolerance curve, although other factors could also account for it independent of insulin resistance. Incorporating insulin resistance in our model and assuming it to work only in the post feeding state, patterns 1, 2, 5, 6, 9, 16 and 17 could be explained but not others. In short, our model does not rule out insulin resistance as a phenomenon, but implies that it may not be central to T2DM. Primary vasculature impairment reducing glucose and insulin transport in mutually correlated or uncorrelated manner could explain all patterns and therefore makes the most parsimonious causal hypothesis. Particularly, assuming that altered vascular function affects *K_1_* early followed by *K_8_* and *K_9_* is sufficient to explain all the patterns without alteration in *I_s_* or any other factor. The apparent β cell dysfunction is an inevitable effect of higher degree of vascular dysfunction and therefore may not be needed as an independent causal factor.

Being open to alternative possibilities is an important virtue of science and it is particularly important with the limited clinical success of prevalent thinking along with mounting anomalous findings. Prevention of T2DM on a global scale has largely failed and treatment has limited and inconsistent success in arresting mortality and morbidities associated with T2DM (Brown et al., 2004; DeFronzo, 2010; Rosengren et al., 2008, Lee et al 2021). Therefore, exploring alternative possible interpretations is a need of the time.

Triggered by multiple anomalies in the theory of glucose homeostasis and the origins of diabetic hyperglycemia, we have articulated here an alternative paradigm which potentially resolves in a logically and mathematically consistent manner many of the anomalous findings. Being mathematically and logically sound and compatible with evidence is not a sufficient proof of a theory but certainly reflects on its potential to develop into a new alternative paradigm. For a large field such as T2DM multiple efforts would be needed to evaluate competing paradigms. We have suggested a few more testable predictions that can help in this task. There can be more possible ways of testing them which should come to light and used to reach robust conclusions which have the potential to change the clinical course of prevention as well as treatment of an important global health problem.

If our proposed causal inference that vascular dysfunction is primary to T2DM is supported by more careful investigations, it explains the failure of glucose normalizing treatment in arresting diabetic complications. Simultaneously it suggests alternative lines of treatments which should focus on normalizing vascular and neuronal function rather than focusing on glucose normalization. The deficiencies of stimuli normally required for growth factors and other angiogenic and neuroprotective factors can be forecasted as the best bet for the new approach. However, rigorous efforts are needed to strengthen the evidence base for selecting the right one amongst the alternative paradigms.

